# Postweaning development influences endogenous VPAC1 modulation of LTP induced by theta-burst stimulation: a link to maturation of the hippocampal GABAergic system?

**DOI:** 10.1101/2024.02.18.580862

**Authors:** M Gil, A Caulino-Rocha, M Bento, NC Rodrigues, A Silva-Cruz, JA Ribeiro, D Cunha-Reis

## Abstract

Long-term potentiation (LTP) induced by theta-burst stimulation (TBS) undergoes postweaning developmental changes partially linked to GABAergic circuit maturation. Endogenous VIP acting on VPAC_1_ receptors strongly influences LTP induced by theta-burst stimulation (TBS), an effect dependent on GABAergic transmission. Although VPAC_1_ receptor levels are developmentally regulated during embryogenesis, its variation along postweaning development is unknow, as is VPAC_1_ modulation of LTP or its relation to hippocampal GABAergic circuit maturation. As such, we investigated how VPAC_1_ modulation of LTP adjusts from weaning to adulthood along with GABAergic circuit maturation. As described, LTP induced by TBS (5×4) (5 bursts, 4 pulses delivered at 100Hz) was increasingly greater from weaning to adulthood. The influence of the VPAC_1_ receptor antagonist PG 97-269 (100nM) on TBS-induced LTP was much larger in *juvenile* (3-week-old) than in *young-adult* (6-7-week-old) or *adult* (12-week-old) rats. This effect was not associated to a developmental decrease in synaptic VPAC_1_ receptor levels. However, an increase in pre and post synaptic GABAergic synaptic markers, suggests an increase in the number of GABAergic synaptic contacts that is more prominent than the one observed in glutamatergic connections during this period. Conversely, endogenous VPAC_2_ receptor activation did not significantly influence of TBS-induced LTP. VPAC_2_ receptor levels enhance pronouncedly during postweaning development, but not at synaptic sites. Given the involvement of VIP interneurons in several aspects of hippocampal-dependent learning, neurodevelopmental disorders, and epilepsy, this could provide important insights into the role of VIP modulation of hippocampal synaptic plasticity during normal and altered brain development potentially contributing to epileptogenesis.

## Introduction

Vasoactive intestinal peptide (VIP), a neuropeptide present exclusively in hippocampal interneurons [1], is a modulator of hippocampal synaptic transmission and excitability[2]. Its endogenous actions are believed to be most relevant in conditions eliciting high frequency firing such as occurring during hippocampal synaptic plasticity and seizures [3,4]. Its actions are mediated by hippocampal VPAC_1_ and VPAC_2_ receptors that integrate the VIP/PACAP family of G protein-coupled receptors [3]. These receptors bind with similar affinity the neuropeptide pituitary adenylate cyclase-activating polypeptide (PACAP), yet another receptor in this family, PAC_1_, has a significantly higher affinity for PACAP than for VIP.

The hippocampal actions of VIP are largely dependent on GABAergic transmission [5–7] and involve multiple cellular and molecular targets, as expected from the expression of VIP in different interneuron populations with different target selectivity [2] and from the opposing actions of VIP on exocytotic GABA release when activating VPAC_1_ or VPAC_2_ receptors [8]. Nevertheless, VIP enhancement of synaptic transmission to CA1 pyramidal cell dendrites is mostly mediated by VPAC_1_ receptors located in the *strata oriens* and *radiatum* [6,9,10]. This, in turn, triggers disinhibition of pyramidal cell dendrites by inhibiting dendritic-targeting GABAergic interneurons [7]. Notwithstanding, VIP also enhances pyramidal cell excitability in the absence of GABAergic transmission [7,11], an effect believed to occur through activation of VPAC_2_ receptors in pyramidal cell bodies [12].

Long-term potentiation (LTP) as induced by theta-burst stimulation (TBS), a form of activity dependent synaptic plasticity, is a key cellular mechanism for memory storage [13,14]. TBS is a sequence of electrical stimuli that mimics CA3 and CA1 pyramidal complex-spike cell discharges perceived during the hippocampal theta rhythm (3-7Hz). This EEG pattern is linked to hippocampal spatial memory formation believed to work as a ‘tag’ for short term memory processing [14,15]. TBS triggers an early LTP, acting through suppression of feedforward inhibition by a priming burst /single pulse. This allows for enough depolarization to trigger NMDA receptors [16,17], through an indirect mechanism that involves activation of GABA_B_ autoreceptors, thus supressing GABA release from feedforward interneurons [18] and synaptic GABA availability.

Endogenous VIP, acting on VPAC_1_ receptors, is an important endogenous modulator of NMDA-dependent TBS-induced LTP as well as LTD and depotentiation in the CA1 area of the hippocampus [4,19]. This is a prospective indirect effect through modulation of disinhibition since it is fully dependent on GABAergic transmission. Yet this, together with previous reports that VIP directly enhances NMDA currents in CA1 pyramidal cells, an effect mimicked by VPAC_2_ and to a lesser extent by VPAC_1_ selective agonists [20], highlights the diversity and importance of VIP in the regulation of hippocampal synaptic plasticity phenomena. Accordingly, VIP is a crucial endogenous modulator of several hippocampal-dependent learning tasks [21–25]. Furthermore, VIP deficient mice fail to develop hippocampal-dependent learning skills like reversal learning [26] while the VIP knockout is inviable. VIP interneuron dysfunction is also implicated in three prominent neurodevelopmental disorders, Rett syndrome, Dravet Syndrome and Down’s syndrome [27–29] and may contribute to enhanced seizure susceptibility and epileptogenesis associated with altered synaptic plasticity in these developmental disorders as in other acquired epilepsies [3,30–32].

The activity of hippocampal GABAergic interneurons influences not only the balance between excitation and inhibition but also a complex network of circuits regulating feedforward, feedback, and disinhibitory mechanisms [33]. Synaptic inhibition at pyramidal cell dendrites is crucial for selective Ca^2+^-dependent input selectivity and precision of LTP induction [34] yet the long lasting plasticity responses of interneurons to mild TBS remain largely unknown. Accumulated evidence suggests that interneurons serve as mediators of normal circuit development by regulating critical period plasticity and shaping the formation of sensory maps [33]. Furthermore, although interneurons are present in the hippocampus from early postnatal development, a few of these, like parvalbumin-expressing interneurons only fully develop and set in human hippocampal circuits until adulthood [35] This can dramatically influence the dynamics of hippocampal inhibition from weaning to adulthood, and its control of synaptic plasticity phenomena. We showed recently that TBS-induced LTP undergoes post-weaning developmental maturation until adulthood [36], and preliminary data from our lab suggests this may be related to postweaning maturation of GABAergic circuits [37].

In this paper we investigated the postweaning developmental changes in VPAC_1_ receptor modulation of hippocampal LTP induced by mild TBS by endogenous VIP and their correlation with developmental changes in the expression of VIP and VIP receptors, GABAergic and glutamatergic pre and postsynaptic markers and overall synaptic density.

## Material and Methods

The experiments were performed in *juvenile* (3 weeks old), *young adult* (6-7 weeks old) and *adult* (12 weeks old) male outbred Wistar rats (Charles River, Barcelona, Spain) essentially as previously described [36,38] and all protocols and procedures were performed according to ARRIVE guidelines for experimental design, analysis, and their reporting. Animal housing and handing was performed in accordance with the Portuguese law (DL 113/2013) and European Community guidelines (86/609/EEC and 63/2010/CE). The animals were anesthetized with fluothane, decapitated, and the right hippocampus dissected free in ice-cold artificial cerebrospinal fluid (aCSF) of the following composition in mM: NaCl 124, KCl 3, NaH_2_PO_4_ 1.25, NaHCO_3_ 26, MgSO_4_ 1.5, CaCl_2_ 2, glucose 10, and gassed with a 95% O_2_ - 5% CO_2_ mixture.

### LTP experiments

Hippocampal slices (400 µm thick) were cut perpendicularly to the long axis of the hippocampus with a McIlwain tissue chopper, then they were kept in a resting chamber in gassed aCSF at room temperature 22°C–25°C for at least 1 h to allow energetic and functional recovery. Each slice was transferred at a time to a submerged recording chamber of 1 ml capacity, and continuously superfused at a rate of 3 ml/min with gassed aCSF at 30.5°C. To obtain electrophysiological recordings slices were stimulated (rectangular pulses of 0.1 ms) through bipolar concentric wire electrodes placed in two separate sets (S1 and S2) of the Schaffer collateral/commissural fibres in the *stratum* radiatum. Responses were evoked alternately on the two pathways every 10 s and thus each pathway was stimulated every 20 s (0.05Hz). Evoked field excitatory post-synaptic potentials (fEPSPs, Fig. 1.A) were recorded extracellularly from CA1 *stratum* radiatum using aCSF-filled micropipettes. The initial stimulus intensity was chosen to elicit a field excitatory post-synaptic potential (fEPSP) of 600–900 mV amplitude (about 50% of the maximal response), while minimizing contamination by the population spike, and of similar magnitude in both pathways. The averages of six consecutive fEPSP responses from each pathway were obtained, measured, graphically plotted and recorded for further analysis with a personal computer using the LTP software [39]. The slope of the initial phase of the potential was used to quantify fEPSPs intensity. At the end of the experiments, the independence of the two pathways was tested by studying paired-pulse facilitation (PPF) across both pathways, less than 10% facilitation being usually observed. PPF was elicited by stimulating the two Schaffer pathways with a 50 ms interval between pulses. The ratio P2/P1 between the fEPSP slopes elicited by the second P2 and the first P1 stimuli were used to quantify synaptic facilitation.

**Figure 1.**
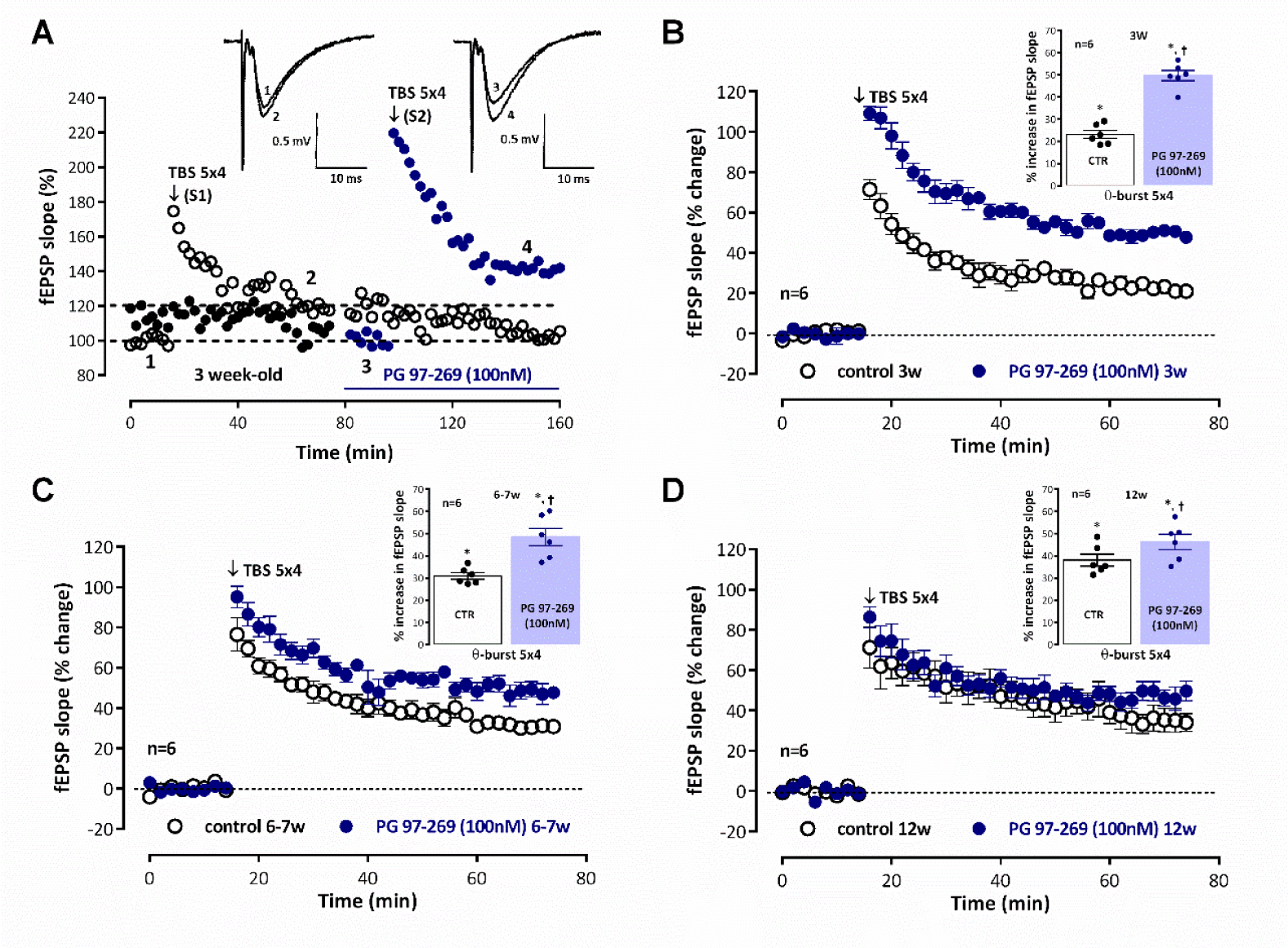
Endogenous inhibition of hippocampal CA1 long-term mild-TBS induced LTP elicited by VIP acting on VPAC_1_ receptors is attenuated from weaning to adulthood. **A.** Time-course of changes in fEPSP slope caused by theta-burst stimulation (5 bursts at 5 Hz, each composed of four pulses at 100 Hz, *mild TBS(5×4)*) in a typical experiment showing both the control pathway (-○-, S1, absence of added drugs) and the test pathway (-●-, S2), to which the VPAC_1_ receptor antagonist PG 97-269 (100nM) was added 30 min before LTP induction, in the same slice obtained from a *juvenile* rat (3 weeks). TBS(5×4) was delivered after obtaining a stable baseline for at least 16 min. ***Inset:*** Traces of fEPSPs obtained in the same experiment before (time point **1**) and 50-60 min after (time point **2**) theta burst stimulation in control conditions and before (time point **3**) and 50-60 min after (time point **4**) theta burst stimulation in the presence of PG 97-269 (100nM). Traces are the average of eight consecutive responses and are composed of the stimulus artifact, the presynaptic volley and the fEPSP. Averaged time-course of changes in fEPSP slope caused by theta-burst stimulation (*mild TBS (5×4))* in the absence (-○-) and in the presence (-●-) of the selective VPAC_1_ receptor antagonist PG 97-269 (100nM) in *juvenile* rats (3-week-old, **B**.), *young-adult* rats (6-7-week-old, **C**.) and *adult* rats (12-week-old, **D**.). Control and test conditions (absence and presence added drugs) were evaluated in independent pathways in the same slice. ***Inset**:* LTP magnitude estimated from the averaged enhancement of fEPSP slope observed 50-60 min, after *mild TBS (5×4)* in the absence of added drugs (-○-) and in the presence of PG 97-269 (-●-,100nM) in *juvenile* rats (**B**.), *young-adult* rats (**C**.) and *adult* rats (**D**.). Values (**B. - D**.) are the mean ± S.E.M. *p < 0.05 (Student’s t test) as compared to the null hypothesis. ⩾p < 0.05 (paired Student’s t test) as compared to the LTP obtained in control conditions (absence of added drugs) for the same age.

When a stable fEPSP slope baseline was observed for at least 20 min, LTP was induced either by *mild TBS* (five trains of 100 Hz, 4 stimuli, separated by 200 ms) or a *moderate TBS* (fifteen trains of 100 Hz, 4 stimuli, separated by 200 ms). The intensity of the stimulus was not changed during these stimulation protocols. LTP was taken as the % change in the average slope of fEPSPs observed from 50 to 60 min after the induction protocol, in relation to one measured during the 10 min that preceded TBS. Control and test conditions were tested in independent pathways in the same slice. In all experiments S1 always refers to the first pathway (left or right, randomly assigned) to which TBS was applied. Test drugs were added to the perfusion solution 20 min before TBS stimulation of the test pathway (S2) and were present until the end of the experiment.

### Western blot analysis of synaptic proteins

The hippocampi of 3 week-old, 6-7-week-old and 12-14 week-old rats were collected in sucrose solution (320mM Sucrose, 1mg/ml BSA, 10mM HEPES e 1mM EDTA, pH 7,4) containing protease (complete, mini, EDTA-free Protease Inhibitor Cocktail, Sigma) and phosphatase (1 mM PMSF, 2 mM Na_3_VO_4_, and 10 mM NaF) inhibitors, homogenized with a Potter-Elvejham apparatus and either total hippocampal membranes or hippocampal synaptosomes were isolated as described [36,40]. For isolation of ***total hippocampal membranes***, the hippocampal homogenates were centrifuged at 1500g for 10 min. The supernatant was collected and further centrifuged at 14000g for 12 min. The pellet was washed twice with modified aCSF (20mM HEPES, 1mM MgCl_2_, 1.2mM NaH_2_PO_4_, 120mM NaCl; 2.7mM KCl, 1.2mM CaCl_2_, 10mM glucose, pH 7.4) also containing protease and phosphatase inhibitors and resuspended in 300µl modified aCSF *per* hippocampus. For isolation of ***hippocampal synaptosomes***, homogenates were centrifuged at 3000 g for 10 min at 4° C; the supernatant was then centrifuged at 14 000 g for 12 min at 4° C and the pellet resuspended in 3 ml of a Percoll 45% (v/v) in modified aCSF. The top layer (synaptosomal fraction) obtained after centrifugation at 14 000 g for 2 min at 4°C, was washed twice with aCSF and resuspended in 200µl modified aCSF *per* hippocampus. Aliquots of these suspensions of hippocampal membranes/synaptosomes were snap-frozen in liquid nitrogen and stored at -80°C until use.

For western blot, samples incubated at 95°C for 5 min with Laemmli buffer (125mM Tris-BASE, 4% SDS, 50% glycerol, 0,02% Bromophenol Blue, 10% β-mercaptoethanol), were run on standard 12% sodium dodecyl sulphate polyacrylamide gel electrophoresis (SDS-PAGE) and transferred to PVDF membranes (pore size 0.45 μm, GE Healthcare Life Sciences). These were then blocked for 1 h with either a 3% BSA solution or 5% milk solution in Tris-buffered saline (20mM Tris, 150mM NaCl) containing 0.1% Tween-20 (TBST), and incubated overnight at 4°C with and either rabbit polyclonal anti-VPAC1 (1:600, Alomone Labs #AVR-001, RRID: AB_2341081), rabbit polyclonal anti-VPAC2 (1:500, Alomone Labs #AVR-002, RRID: RRID: AB_2341082), rabbit polyclonal anti VIP (1:300, Proteintech, # 16233-1-AP, RRID: AB_2878233), mouse monoclonal anti-gephyrin (1:3000, #147011, Synaptic Systems, AB_2810214), rabbit polyclonal anti-PSD-95 (1:750, #CST-2507, Cell Signalling Tech., AB_561221), rabbit polyclonal anti-synaptophysin (1:7500, Synaptic Systems #101002, RRID:AB_887905), rabbit polyclonal anti-VGAT (1:2500, Synaptic Systems #131002, RRID: AB_887871), rabbit polyclonal anti-VGlut1 (1:3000, Synaptic Systems #135302, RRID: AB_887877), and either mouse monoclonal anti-β-actin (1:5000, Proteintech #60008-1-Ig, RRID: AB_2289225) or rabbit polyclonal anti-alpha-tubulin (1:5000, Proteintech #11224-1-AP; RRID: AB_2210206) primary antibodies. After washing the membranes were incubated for 1h with anti-rabbit or anti-mouse IgG secondary antibody both conjugated with horseradish peroxidase (HRP) (Proteintech) at room temperature. HRP activity was visualized by enhanced chemiluminescence with Clarity ECL Western Blotting Detection System (Bio-Rad). Intensity of the bands was evaluated with the Image J software. alpha-tubulin band density was used as a loading control.

## Materials

PG 97-269 and PG 99-465, (Phoenix peptides, Europe) were made up in 0.1mM stock solution in CH_3_COOH 1% (v v-1). The maximal CH_3_COOH concentration added to the slices, 0.001% (v/v) was devoid of effects on fEPSP slope (n=4). Aliquots of the stock solutions were kept frozen at -20°C until use. In each experiment, one aliquot was thawed and diluted in aCSF.

### Statistics

LTP values are depicted as the mean ± S.E.M of n experiments. Each *n* represents a single experiment performed in slices obtained from one different animal for LTP experiments and experiments performed in duplicate for each single animal in western-blot studies. The significance of the differences between the means was calculated using the paired Student’s t-test when comparing control and test conditions for LTP expression in electrophysiological studies, or with repeated measures ANOVA with Tukey’s post-hoc test (when F was significant) when comparing the levels of synaptic proteins in different age groups. P values of 0.05 or less were considered to represent statistically significant differences.

## Results

Recordings of fEPSP obtained under basal stimulation conditions (40-60% of the maximal response in each slice) in hippocampal slices from *juvenile* (3-week-old) rats (Fig. 1.A, raw data from a single experiment) showed an average slope of 0.579±0.036 mV/ms (n=12). When *mild TBS (5×4 pulses with a 200ms interval)* was applied to the control pathway (S1) an LTP was induced, corresponding to a 25.6±1.9% increase in fEPSP slope (n=12, P<0.05) observed 50-60min after TBS. Application of a second *mild TBS* train in the test pathway (S2) in the absence of drugs always resulted in an LTP of similar magnitude (% increase in fEPSP slope of 26.1±3.4%, n=4) to the one observed in the control pathway (S1), i.e., LTP obtained under these experimental conditions was similar in S1 and S2. When *mild TBS* of the test pathway (S2) was delivered in the presence of the VPAC_1_ receptor antagonist PG 97-269 (100nM) LTP was almost twice as large, since the enhancement of fEPSP slope observed 1h latter was of 49.6±2.3%, (n=6), suggesting a strong inhibition of LTP by endogenous VIP acting on VPAC_1_ receptors. This effect was larger than the one previously observed in *young adult* rats (6-7 weeks old) [19]. In this study, fEPSPs in *young adult* rats had an average slope of 0.553±0.022 mV/ms (n=17). Induction of LTP with a *mild TBS* caused a larger potentiation of the fEPSP slope (% increase in fEPSP slope 30.0±1.2%, n=12, P<0.05, Fig 1.C) than the one observed in *juvenile* rats as previously described [36]. When *mild TBS* was delivered to S2 by in the presence of the VPAC_1_ receptor antagonist PG 97-269 (100nM) this resulted in an enhancement of LTP (% increase in fEPSP slope of 48.4±3.9%, n=6, Fig. 1.C) that was less pronounced than the one observed in *juvenile* rats (Fig 1.B). In *adult* rats (12weeks old) the average fEPSP slope was of 0.582±0.017 mV/ms (n=16). The same *mild TBS* paradigm caused an even larger LTP since *mild TBS* now induced a potentiation of the fEPSP slope of 38.1±2.7%, n=6, P<0.05, Fig 1.D). When LTP was induced in the presence of the VPAC_1_ receptor antagonist PG 97-269 (100nM) the enhancement of LTP was even more reduced, since *mild TBS* induced an enhancement of 46.3±3.4% (n=6, Fig. 1D) in fEPSP slope. PG 97-269 (100nM) did not significantly change basal synaptic transmission when added to the hippocampal slices.

Our previous studies did not show any influence of endogenous VIP on LTP induced by *mild TBS* through activation of VPAC_2_ receptors in *young adult* rats[19]. However, there could be postweaning developmental changes in this respect. As shown in Fig. 2, when LTP was induced by *mild TBS* in the presence of the selective VPAC_2_ receptor antagonist PG 99-465 (100nM) there was a mild but significant enhancement of LTP in *juvenile* rats for which an enhancement of 34.2±3.4% (n=5, Fig. 2.A) in fEPSP was observed upon mild TBS stimulation. An even milder tendency to an increase in LTP by VPAC_2_ receptor blockage was observed in *young adult* and *adult* rats, but it was not consistently observed in all animals (Fig. 2.B and 2.C), and this effect did not reach statistical significance. When added to the hippocampal slices PG 99-465 (100nM) did not significantly change basal synaptic transmission.

**Figure 2.**
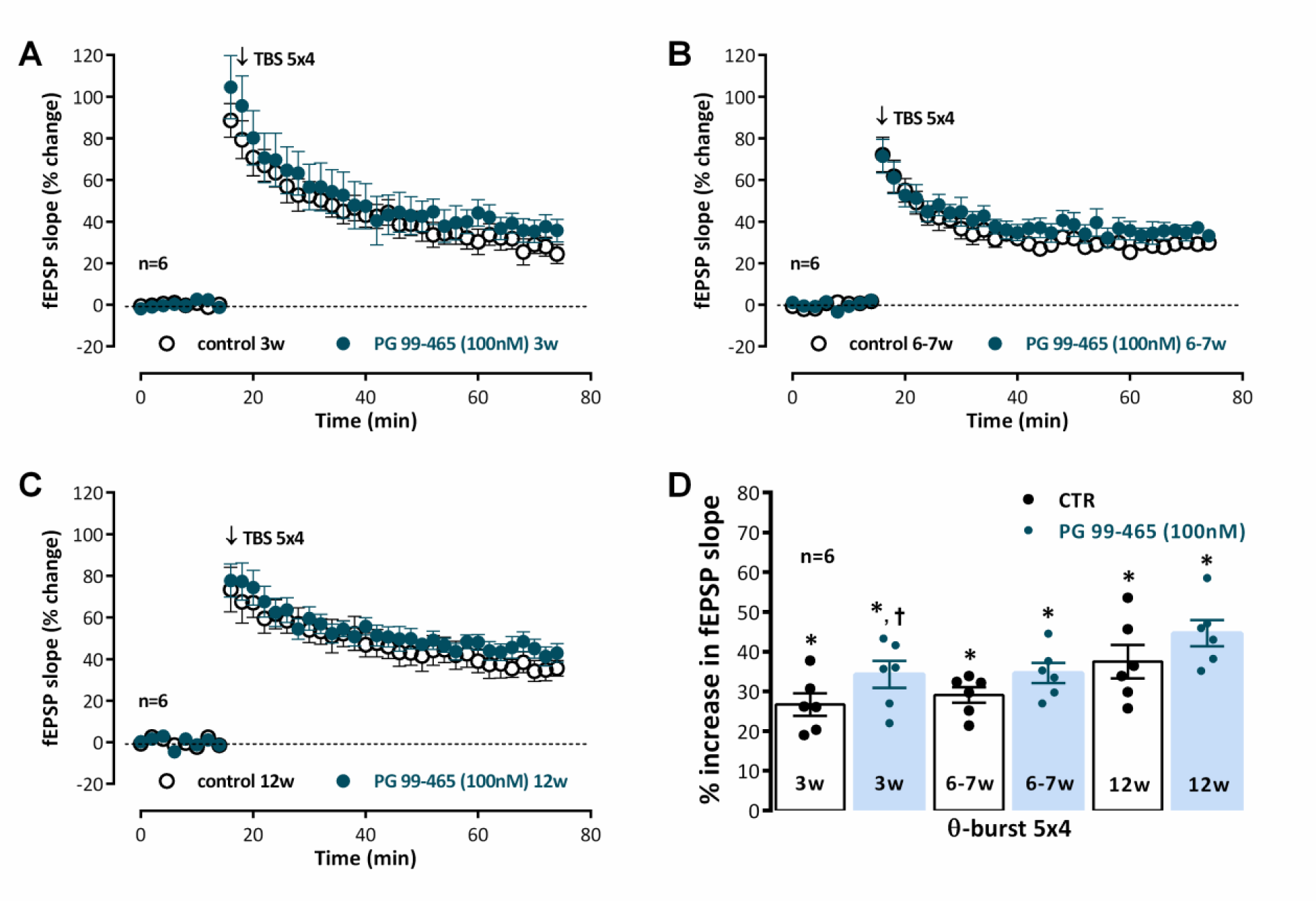
Activation of VPAC2 receptors is not involved in the endogenous control of hippocampal CA1 long-term potentiation of synaptic transmission by VIP from weaning to adulthood. Averaged time-course of changes in fEPSP slope caused by theta-burst stimulation (*mild TBS (5×4))* in the absence (-○-) and in the presence (-●-) of the selective VPAC_2_ receptor antagonist PG 99-459 (100nM) in *juvenile* rats (3-week-old, **A.**), *young-adult* rats (6-7-week-old, **B.**) and *adult* rats (12-week-old, **C.**). Control and test conditions (absence and presence added drugs) were evaluated in independent pathways in the same slice. **D.** LTP magnitude estimated from the averaged enhancement of fEPSP slope observed 50-60 min, after *mild TBS (5×4)* in the absence of added drugs (-○-) and in the presence of PG 97-269 (-●-,100nM) in *juvenile* rats (3w), *young-adult* rats (6-7w) and *adult* rats (12w). Values (**A.** - **D.**) are the mean ± S.E.M. *p < 0.05 (Student’s t test) as compared to the null hypothesis. ⩾p < 0.05 (paired Student’s t test) as compared to the LTP obtained in control conditions (absence of added drugs) for the same age.

Increasing the number of bursts to 15 (*moderate TBS*) enhanced the resulting potentiation evaluated 50-60min after TBS, leading to a 46.2±1.8% (n=5, P<0.05) enhancement in fEPSP slope (Fig 3.A) in *young adult* rats and to a 63.6±3.3% (n=5, P<0.05) enhancement in fEPSP slope (Fig 3.B) in *adult* rats 50-60m after TBS stimulation, as previously described [36]. Using this stronger LTP induction paradigm the influence of endogenous VIP, acting on VPAC_1_ receptors, on LTP expression was altered since the potentiation induced in fEPSP slope by moderate TBS in the presence of the selective VPAC_1_ receptor antagonist PG 97-269 (100nM) was of 59.9±3.1% (n=5, P<0.05, Fig 3.A) in *young adult* rats and of a 56.5±4.4% (n=5, P<0.05, Fig 3.B) in *adult* rats, thus displaying a much lower potentiation of LTP by the VPAC_1_ receptor antagonist in *young adult* rats and a mild but non-significant reversal of the effect in *adult* rats, suggesting that a developmental reshaping of GABAergic synapses, VIPergic interneurons or its activation pathways may be occurring or that VIP receptor expression and its role in hippocampal GABAergic synapses may be altered.

**Figure 3.**
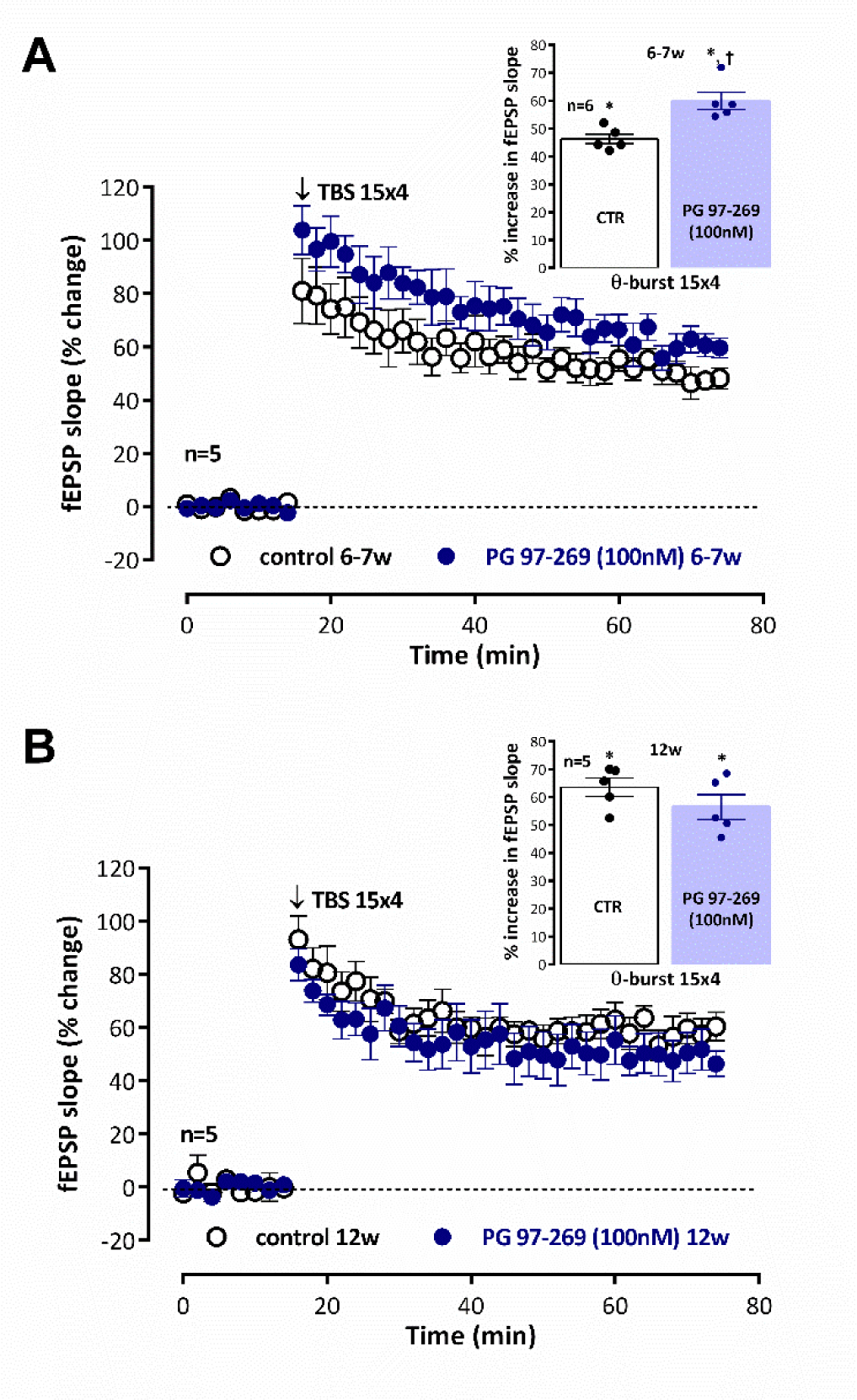
Endogenous inhibition of hippocampal CA1 strong-TBS induced LTP elicited by VIP acting on VPAC_1_ receptors is abolished reaching adulthood. Averaged time-course of changes in fEPSP slope caused by theta-burst stimulation (*mild TBS (5×4))* in the absence (-○-) and in the presence (-●-) of the selective VPAC_1_ receptor antagonist PG 97-269 (100nM) in *young-adult* rats (6-7-week-old, **A.**) and *adult* rats (12-week-old, **B.**). Control and test conditions (absence and presence added drugs) were evaluated in independent pathways in the same slice. *Inset:* LTP magnitude estimated from the averaged enhancement of fEPSP slope observed 50-60 min, after *mild TBS (5×4)* in the absence of added drugs (-○-) and in the presence of PG 97-269 (-●-,100nM) in *young-adult* rats (**A.**) and *adult* rats (**B.**). Values (**A.** - **B.**) are the mean ± S.E.M. *p < 0.05 (Student’s t test) as compared to the null hypothesis. ⩾p < 0.05 (paired Student’s t test) as compared to the LTP obtained in control conditions (absence of added drugs) for the same age.

To elucidate the changes in VIP, VIP receptors, GABAergic and glutamatergic synapses during postweaning development in the hippocampus we studied the levels of VIP VPAC_1_ and VPAC_2_ receptors and synaptic glutamatergic and GABAergic markers by western blot in total hippocampal membranes. The synaptic changes in glutamatergic and GABAergic transmission were inferred from the evolution of GABAergic and glutamatergic pre and postsynaptic markers and its relation to broad-spectrum presynaptic marker synaptophysin, that was taken as an estimator of the overall synaptic density in total hippocampal membranes. The levels of the synaptic vesicle integral protein synaptophysin enhanced from weaning to adulthood by 63.9±14.9% (n=5, P<0.05, Fig 4.I). The levels of both pre- and post-synaptic glutamatergic synaptic proteins VGlutT-1 and PSD-95 were enhanced by 29.9±6.6% (n=5, P<0.05, Fig 4.A and C) and 44.5±11.0% (n=5, P<0.05, Fig 4.A and B), respectively, in relation to the observed protein levels at 3 weeks of age. The increase in pre- and post-synaptic GABAergic synaptic proteins VGAT and gephyrin was more pronounced since VGAT and gephyrin levels enhanced by 71.1±16.0% (n=5, P<0.05, Fig 4.E and G) and 55.4±10.9% (n=5, P<0.05, Fig 4.E and F), respectively, in relation to the protein levels observed at 3 weeks, the age of weaning. By plotting the VGlut1/PSD-95 and VGAT/gephyrin ratios (Fig 4.D and H) we could observe that the ratios do not significantly change along postweaning development, suggesting that the increase in these synaptic markers is indeed representative of an increase in the number of synapses. To estimate the rate of change of glutamatergic and GABAergic synapses vs global synaptic changes in hippocampal tissue we calculated the VGlut1/synaptophysin and the VGAT/synaptophysin immunoreactivity ratios (Fig. 4.K and L). The decrease in the hippocampal VGlut1/synaptophysin ratio from 3 to 12 weeks suggests that glutamatergic synapses do not accompany the global increase in new synapses while GABAergic synapses seem to contribute to this increase in hippocampal synaptic content, as estimated by a stable VGAT/synaptophysin ratio from 3 to 12 weeks.

**Figure 4.**
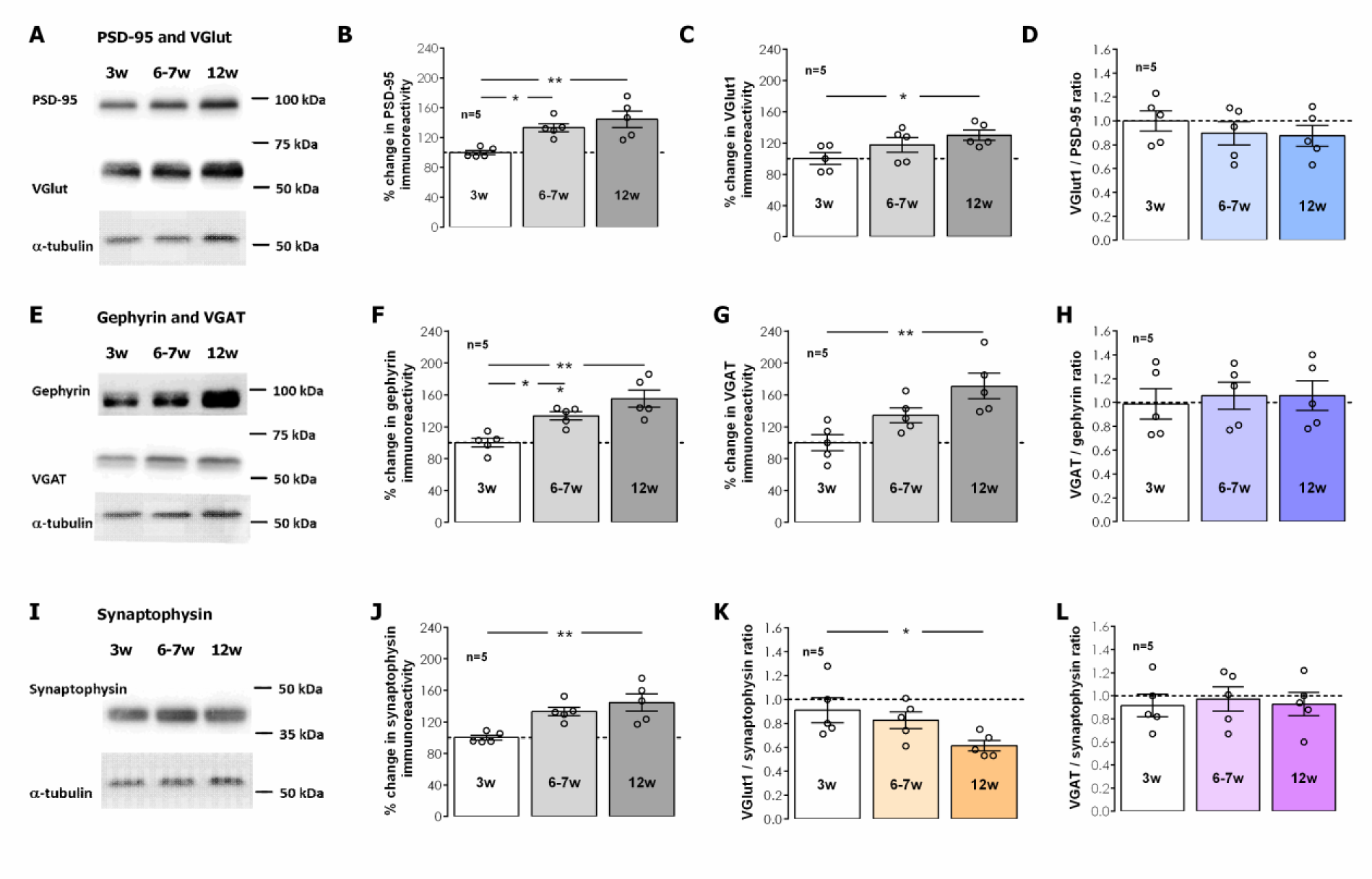
Impact of post-weaning development into adulthood on the levels of structural, GABAergic, and glutamatergic synaptic proteins. **A., E**., and **I.** Western-blot immunodetection of PSD-95, VGlut1, Gephyrin, VGAT and synaptophysin obtained in one individual experiment in which total hippocampal membranes were subjected to WB analysis. Averaged total PSD-95 (**B.**), VGlut1 (**C.**), Gephyrin (**F.**), VGAT (**G.**) and synaptophysin (**J.**) immunoreactivities. VGlut1/PSD-95 (**D.**) ratio and VGAT/Gephyrin ratio (**H.**) for each animal were also plotted as an estimate of synaptic growth. VGlut1/synaptophysin (**K.**) ratio and VGAT/Synaptophysin ratio (**L.**) are also depicted and were used as an estimate of GABAergic vs glutamatergic synaptic enhancement. Values are mean ± S.E.M of five independent experiments performed in duplicate and were normalized to the immunoreactivity of α-tubulin, used as loading control. 100% - averaged target immunoreactivity obtained for 3-week-old animals. **\*** Represents p < 0.05 **\*\*** represents p < 0.01 (*ANOVA*, Tukey’s multiple comparison test) as compared to protein levels in 3-week-old animals or as otherwise indicated.

In total hippocampal membranes, VPAC_1_ receptor immunostaining was detected by two independent bands located at 55 and 75KDa, respectively, likely reflecting two different N-glycosylation levels, a feature that is also relevant for VPAC_1_ membrane targeting, as previously described [41]. The levels of both VPAC_1_ and VPAC_2_ receptors in total hippocampal membranes displayed a gradual postweaning developmental enhancement, that reached 42.5±8.9% enhancement in 12-week-old rats for VPAC_1_ receptors (Fig. 5.A-B, n=5, P<0.05) and was even more pronounced for VPAC_2_ receptors, for which the receptor levels nearly doubled (% increase 93.5±15.7, Fig. 5.D-E, n=5, P<0.05) in 12-week-old rats when compared to 3-week-old animals. By plotting the VPAC_1_/ and VPAC_2_/synaptophysin ratios (Fig 5.C and F) we were able to infer that the increase in VPAC_1_ receptors accompanied the overall gain in synaptophysin yet VPAC_2_ receptors were increasing even further, suggesting this increase was occurring also in non-synaptic locations.

**Figure 5.**
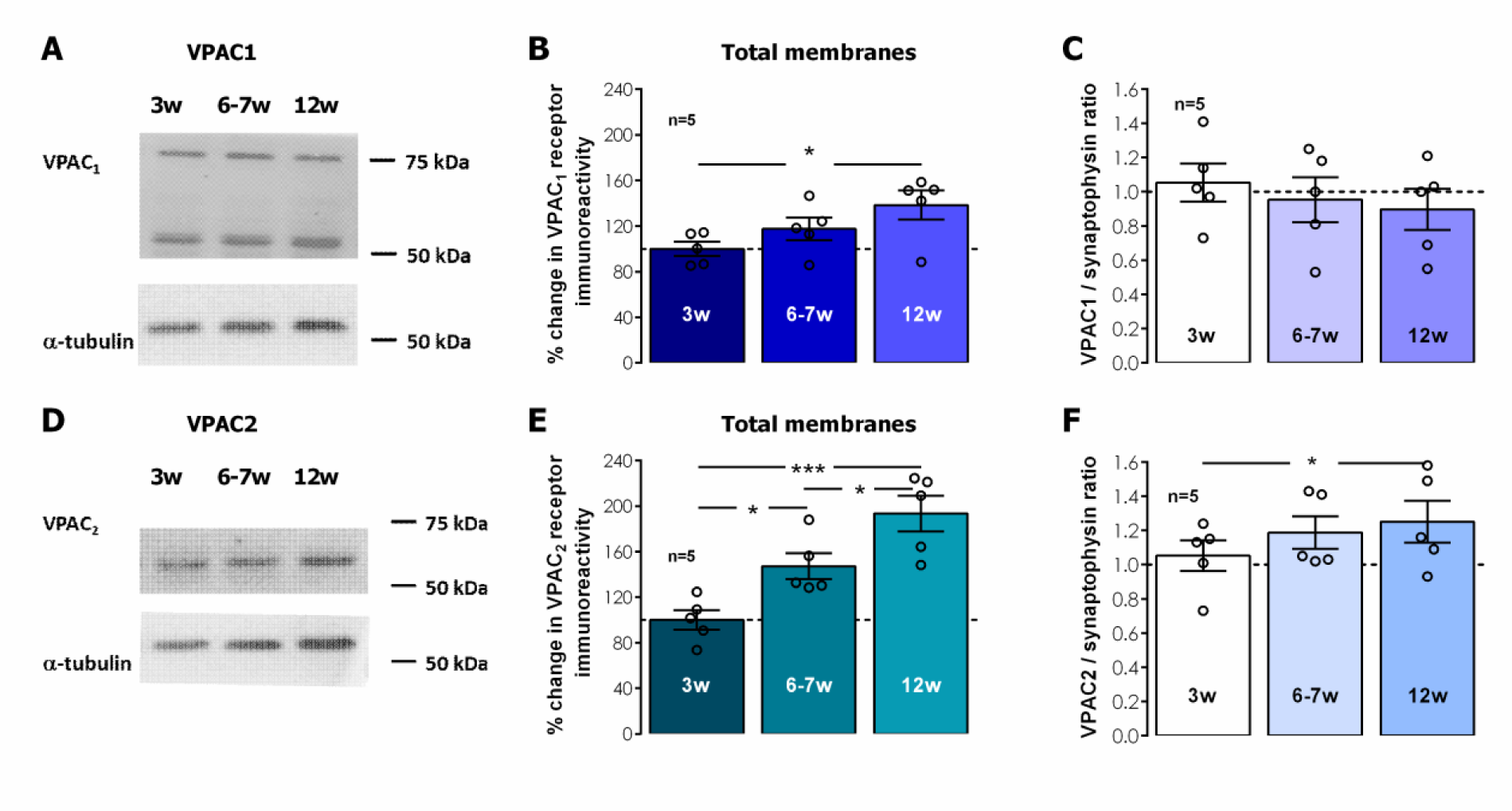
Influence of post-weaning development into adulthood on the global hippocampal levels VPAC_1_ and VPAC_2_ receptors. **A.** and **D.** Western-blot immunodetection of VIP VPAC_1_ and VPAC_2_ receptors obtained in one individual experiment in which hippocampal synaptosomes were subjected to WB analysis. Averaged total VPAC_1_ (**B.**) and VPAC_2_ (**C.**) immunoreactivity. Values are mean ± S.E.M of five independent experiments performed in duplicate and were normalized to the immunoreactivity of α-tubulin, used as loading control. 100% - averaged VPAC_1_ or VPAC_2_ immunoreactivity obtained for 3-week-old animals. VPAC_1_/synaptophysin (**K.**) ratio and VPAC_2_/Synaptophysin ratio (**L.**) are also depicted. **\*** Represents p < 0.05 and **\*\*\*** represents p < 0.001 (*ANOVA*, Tukey’s multiple comparison test) as compared to protein levels in 3-week-old animals or as otherwise indicated.

To further clarify the cellular location of these receptors and their relevance for synaptic transmission and synaptic plasticity we performed also western blot experiments to probe for VIP, VPAC_1_ and VPAC_2_ receptors in hippocampal synaptosomes, a preparation that is composed mainly of presynaptic nerve terminals, that often maintain attached part of the postsynaptic density [42]. In hippocampal synaptosomes, both VIP and to a lesser extent VPAC_1_ receptors increased along post-weaning development by 58.3±15.9%and 43.7±10.1% (n=5, P<0.05, Fig 6.A-B and E-F) at 12 weeks of age, respectively, suggesting that VIP and VPAC_1_ containing nerve terminals enhance substantially along postweaning development. Conversely, VPAC_2_ receptors did not significantly change hippocampal synaptosomes from 3-to 12-week-old rats (% change at 12 weeks vs. 3 weeks 20.3±11.7%, n=5, P>0.05, Fig 6.C-D).

**Figure 6.**
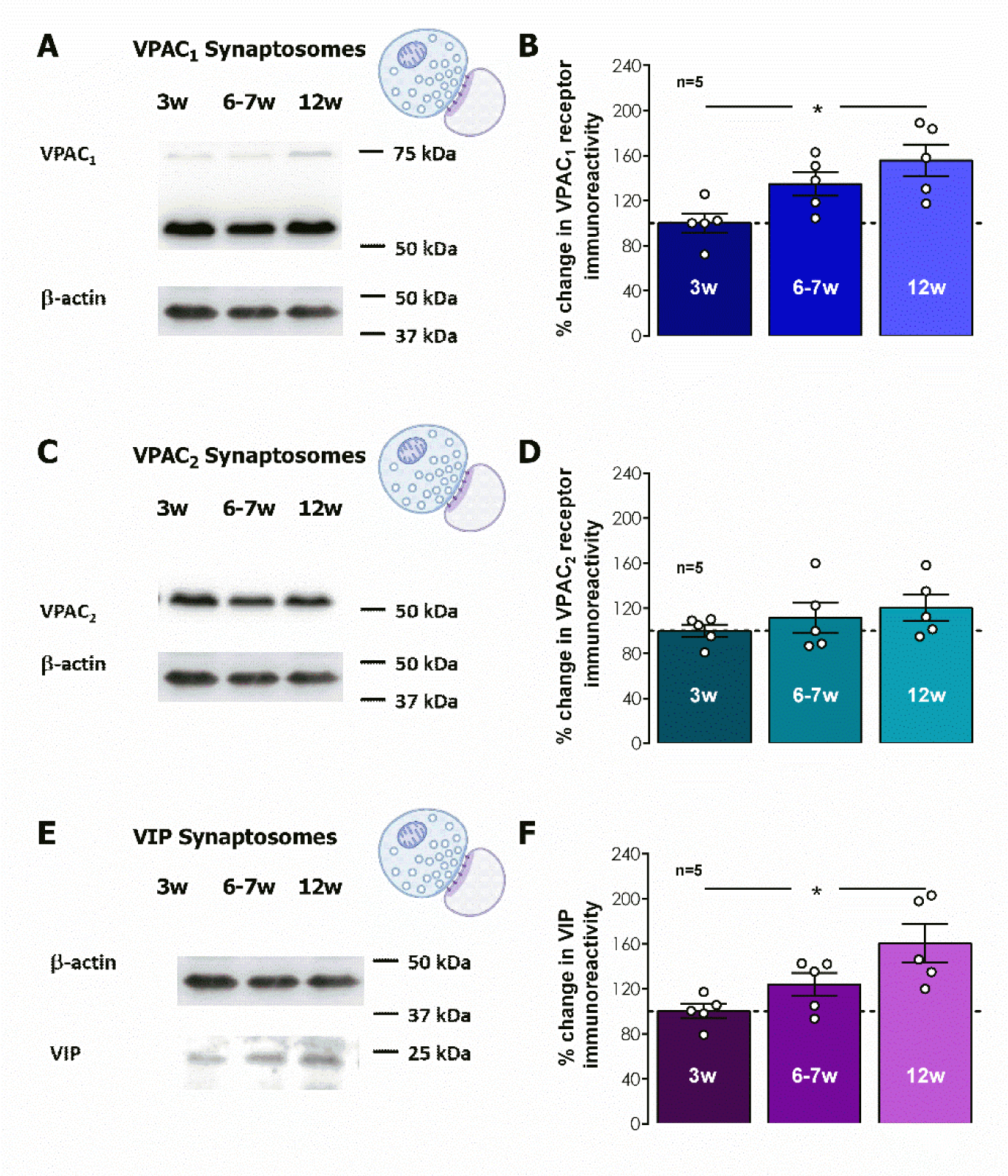
Influence of post-weaning development into adulthood on the on VIP synaptic content and synaptic VPAC_1_ and VPAC_2_ receptors. **A., C.**, and **E.** Western-blot immunodetection of VIP, VPAC_1_ and VPAC_2_ receptors obtained in one individual experiment in which hippocampal synaptosomes were subjected to WB analysis. Averaged total VPAC_1_ (**B.**), VPAC_2_ (**D.**) and VIP (**F.**) immunoreactivities. Values are mean ± S.E.M of five independent experiments performed in duplicate and were normalized to the immunoreactivity of α-tubulin, used as loading control. 100% - averaged VIP, VPAC_1_ or VPAC_2_ immunoreactivity obtained for 3-week-old animals. **\*** Represents p < 0.05 (*ANOVA*, Tukey’s multiple comparison test) as compared to protein levels in 3-week-old animals.

## Discussion

The main findings of the present work are that: 1) VPAC_1_ receptor-mediated modulation of LTP induced by *mild TBS* in the hippocampal CA1 area by endogenous VIP is progressively weaker from weaning (3 week-old) through adulthood (12-week-old) in Wistar rats; 2) A stronger LTP-inducing train (*moderate TBS*) elicited an increased magnitude and stability of LTP in *young-adult* (6-7-week-old) and *adult* (12-week-old) Wistar rats, that was even less influenced by endogenous VIP acting on VPAC_1_ receptors ; 3) The LTP elicited by *mild TBS* was only weakly dependent on endogenous VPAC_2_ receptor activation in *juvenile* (3-week-old) rats; 4) These observations were concurrent with an increase in both GABAergic and glutamatergic pre- and post-synaptic markers in total hippocampal membranes from weaning to adulthood, the enhancement in GABAergic being more prominent; 5) There was an increase in both VPAC_1_ and VPAC_2_ receptor levels in total hippocampal membranes during this developmental period while 6) VIP levels enhanced markedly, while only VPAC_1_, but not VPAC_2_, receptor levels were mildly enhanced in *adult* (12-week-old) vs *juvenile* (3-week-old) rats.

In the present study *mild TBS* (5 bursts, 4 pulses delivered at 100Hz), an LTP-inducing stimulation pattern, elicited a long-lasting potentiation of fEPSP slope that was increasingly larger in 3-, 7-, and 12-week-old rats, as previously described by our group [36]. This is consistent with previous studies showing that during postnatal development there is a reinforcement of the cellular mechanisms leading to LTP expression and stability [14,43,44]. Stronger TBS, achieved by increasing the number of bursts to 15 (*moderate TBS*), enhanced proportionally the resulting potentiation for all age groups studied [36]. *Mild TBS* elicits an LTP that is fully dependent on NMDA and partially dependent on GABA_B_ receptor activation for its induction in our model [36], and is of ideal magnitude for pharmacological studies aiming to improve LTP outcome, while *moderate TBS* elicits a near-maximal LTP for this stimulation pattern [14]. Suppression of feed-forward phasic inhibition, mediated by GABA_B_ autoreceptor inhibition of GABA release, underlies induction of LTP by TBS in excitatory synapses of the hippocampus [14,45,46]. This allows for temporal summation of excitation and sustained depolarization ultimately activating NMDA receptors, a crucial mechanism for TBS LTP induction.

We previously described that LTP induced by mild TBS induced LTP is inhibited by endogenous VIP acting on VPAC_1_ receptors in young adult rats[19]. This work further describes that VPAC_1_ receptor blockade with PG 97-269 elicited an enhancement of LTP and that effect was progressively smaller from weaning, at 3 weeks of age, to adulthood, at 12 weeks of age. In addition, the LTP obtained in the presence of PG 97-269 was very similar for all age groups (46.3 – 49.6%) while LTP induced by *moderate TBS*, an LTP that is near maximal for this stimulation pattern (56.5-59.9%) [14] and could not be further enhanced by VPAC_1_ receptor blockade. Last, the levels of VPAC_1_ receptors increased in both total hippocampal membranes and hippocampal synaptosomes increased from weaning to adulthood. Altogether, this suggests that the decrease in endogenous VIP control of hippocampal CA1 LTP is not due to changes in VPAC_1_ receptor levels and that enhanced LTP afforded by VPAC_1_ receptor blockade reaches a near maximum value for *mild TBS* induced LTP. This is also consistent with previous observations of a developmental reinforcement of the cellular mechanisms leading to LTP expression and stability [14,43,44].

Conversely, as previously described in young-adult rats [19], VPAC_2_ receptor blockade with PG 99-465 did not change LTP of synaptic transmission induced by *mild TBS* in the CA1 area of the hippocampus except for a mild increase in *juvenile* rats, suggesting that endogenous VIP controlling CA1 LTP does not significantly activate VPAC_2_ receptors. This does not preclude an important role for VPAC_2_ receptors in the control of other hippocampal plasticity of neuronal excitability mechanisms, such as modulation of pyramidal cell excitability, as inferred from the main localization of these receptors at the *stratum pyramidale* in the CA1 [9,12,20]. Nevertheless, the enhancement in VPAC_2_ receptor levels observed from weaning to adulthood in total hippocampal membranes was not reproduced in hippocampal synaptosomes, suggesting that it may be occurring either in non-neuronal cells, or away from synaptic sites. Astrocytes are important players in neuronal communication and express functional VPAC_2_ receptors regulating glutamate uptake [47,48], suggesting a major influence in the control of hippocampal synaptic activity and plasticity by endogenous VIP. This should be further investigated but is beyond the scope of the current paper.

Feed-forward phasic inhibition to CA1 pyramidal cells is mediated by physiologically and morphologically distinct GABAergic interneurons[49,50] and its *TBS-induced* suppression, mediated by GABA_B_ autoreceptors, is important, but not fundamental, for TBS-induced CA1 LTP [18,36], as *mild* TBS also activates postsynaptic mechanisms leading to a sustained potentiation of fast GABA_A_ synaptic transmission to pyramidal cells [51]. The net result of all these mechanisms is a potentiation of synaptic transmission by *mild TBS*, and blocking GABA_A_ receptors using bicuculline elicits an enhancement of LTP in the CA1 area [36,52]. Altogether, this suggests that further excitatory and inhibitory pathways, like transient changes in tonic inhibition during LTP induction may determine the outcome of LTP induced by TBS [14,53]. This is corroborated by observations that, both synaptic and non-synaptic inhibition can hinder spike backpropagation and Ca^2+^ spikes in CA1 pyramidal cell dendrites [34,53], an essential mechanism in TBS-induced LTP.

In this study we observed a developmental decrease in VPAC_1_ receptor influence on hippocampal LTP, even though there was a small but significant synaptic enhancement in both VIP and VPAC_1_ receptors, which could be considered a paradox. Furthermore, VPAC_1_ receptors mediate modulation of hippocampal synaptic transmission and plasticity through mechanisms that are fully dependent on GABAergic transmission [6,19,54], VPAC_1_ receptors are important nerve terminal modulators of GABA release [8] and VIP is present in the hippocampus exclusively in three distinct interneuron populations with very distinct target selectivity [1,55]. In view of all this, and of the fact that developmental changes in tonic GABAergic inhibition at pyramidal cell dendrites were associated with developmental changes in LTP expression [56], with relevance for LTP induced by *mild* TBS at developmental stages in this study, the distinct capacity of the VPAC_1_ receptor antagonist (and thus endogenous VIP) to influence the outcome of LTP induced by mild TBS may be conditioned by postweaning maturation of GABAergic circuits. In fact, we previously showed that VPAC_1_ receptor mediated inhibition of CA1 LTP by endogenous VIP is majorly occurring by curtailing disinhibition impinging on phasic feed-forward inhibition [19]. As such, a developmental enhancement in the influence of tonic inhibition on LTP expression would reduce the influence of endogenous VIP acting on VPAC_1_ receptors to modulate the same LTP.

To further explore the possible influence of GABAergic system maturation during postweaning development on modulation of hippocampal LTP in our rat model, we also investigated the alterations in GABAergic and glutamatergic synapses. Changes in synaptic GABAergic and glutamatergic synaptic markers were studied in total hippocampal membranes, which allowed us to perceive the increase in the total number of synapses in the whole tissue, while analysis of the ratios between pre- and post-synaptic GABAergic or glutamatergic markers allowed us to ascribe these changes to synaptic rather than single molecule alterations. In addition, synaptophysin levels were taken as a measure of the global increase in the number of synapses. With this strategy we were able to demonstrate that the increase in the number of hippocampal GABAergic synapses is larger than the increase in the number of glutamatergic synapses along this postweaning developmental period. Studies on the maturation of hippocampal GABAergic transmission and its relations to GABAergic synapse establishment and synaptic plasticity are more often focused on early preweaning brain development [57], but accumulated evidence suggests that maturation of hippocampal circuits continues to progress into childhood and adolescence [58]. This involves not only maturation of resting properties and the setting up of new or altered synaptic connections but also the migration and insertion of new neurons, and involved most prominently interneurons and their synaptic spines [35,58,59]. The observations in the present work are consistent with these studies.

The findings in this paper are of particular interest for the understanding of alterations of hippocampal synaptic plasticity in several developmental brain pathologies that have concomitantly been associated with alterations of VIP or VIP immunoreactive interneurons [33,60,61]. VIP and VPAC_1_ receptors are upregulated in mouse models of Down syndrome [61], a disease associated with excessive GABAergic inhibition, which also displays impaired synaptic plasticity specifically in TBS but not high frequency stimulation-induced LTP *in vitro*[62]. *In vivo*, the deficits in LTP and learning and memory deficits in this model can be rescued by selective GABA antagonists and an α5-selective GABA_A_ inverse agonist [63,64], a subunit associated with both tonic and synaptic inhibition by GABA_A_ receptors in the hippocampus [65]. Enhanced VIP and VPAC_1_ receptor levels may also be involved in restraining disinhibition contributing to impaired LTP in this model. Furthermore, VIP interneuron impairment promotes *in vivo* circuit dysfunction in the cerebral cortex of and autism-related behaviors in Dravet syndrome, a disease caused by mutation of the gene coding for voltage-gated sodium channel subunit Nav1.1 and Rett syndrome, associated with mutations in the gene coding for methyl-CpG binding protein 2 (MECP2) located on the X chromosome, and both associated with recurrent seizures that ultimately lead to epilepsy [33]. Although the firing patterns of irregular spiking VIP expressing interneurons were altered in a mouse model of Dravet syndrome (Scn 1a^+/-^) [27], suggesting this could contribute to Dravet syndrome cognitive pathogenesis. The alterations in the hippocampus are likely secondary to the development of seizures, but should be further investigated, especially in what regards the development of epilepsy, since it has been recently demonstrated that inhibition of VIP-expressing interneurons of the ventral subiculum was sufficient to reduce seizures in the intrahippocampal kainic acid model of epilepsy, thus suggesting a prominent role of these cells in seizure propagation and epileptogenesis [32].

In conclusion, this paper reveals the postweaning alterations in VPAC_1_ receptor modulation of TBS-induced LTP by endogenous VIP in the rat hippocampus. Besides providing us with important insights into VPAC_1_ receptors as putative drug targets aiming to improve synaptic plasticity, and ultimately memory, this study also sheds light into the ongoing physiological adaptations GABAergic transmission and in the control of synaptic plasticity during postweaning development, a developmental period that is often overlooked in brain development. Understanding these physiological adaptations may prove useful to also understand the pathophysiology of neurodevelopmental disorders affecting the hippocampus thus providing a better guide to the discovery of new therapeutic targets.

## Data accessibility statement

The data that support the findings of this study are readily available on request from the corresponding author. The data are not publicly available due to privacy or ethical restrictions.

## Ethics statement

All protocols and procedures were performed according to ARRIVE guidelines for experimental design, analysis, and their reporting. Animal housing and handing was performed in accordance with the Portuguese law (DL 113/2013) and European Community guidelines (86/609/EEC and 63/2010/CE). Experimental protocols were approved by the Ethics Committee of the Faculty of Medicine University of Lisbon (Comissão de ética para a saúde do CHLN/FMUL).

## Conflict of interests

The authors have no conflict of interests to the publication of this paper.

## Author contribution

***M Gil:*** formal analysis and methodology; ***A Caulino-Rocha:*** formal analysis and methodology; ***M Bento:*** formal analysis and methodology; ***NC Rodrigues:*** formal analysis and methodology; ***A Silva-Cruz:*** formal analysis and methodology; ***JA Ribeiro:*** resources, supervision, and writing – review and editing and ***D Cunha-Reis:*** formal analysis and methodology, resources, supervision, funding acquisition, project administration, and writing – original draft, review, and editing.

## Funding

Work was supported by national and international funding managed by Fundação para a Ciência e a Tecnologia (FCT, IP), Portugal. **Grants:** UIDB/04046/2020 (DOI: 10.54499/UIDB/04046/2020) and UIDP/04046/2020 (DOI: 10.54499/UIDP/04046/2020) Centre grants to BioISI, and research grants PTDC/SAU-NEU/103639/2008 and FCT/POCTI (PTDC/SAUPUB/28311/2017) EPIRaft grant (to DC-R). **Fellowships:** SFRH/BPD/81358/2011 to DCR and **Researcher contract:** Norma Transitória - DL57/2016/CP1479/CT0044 to DCR (DOI: 10.54499/DL57/2016/CP1479/CT0044). Marta Gil was in receipt of an Erasmus+ Mobility Fellowship from the Faculty of Biological Sciences, University of Wrocław. Funding sources made no contribution to the writing, research plan and decision to publish this paper.

## Acknowledgements

We acknowledge the Institute of Physiology, FMUL, for animal housing facilities and Raquel Batista Dias for technical contribution with initial LTP experiments in juvenile rats.

## Notes

### Competing Interest Statement

The authors have declared no competing interest.

